# Melatonin Protects T-2 toxin-induced neuronal stress through Acetylcholinesterase & Cytochrome P450 receptor-mediated signaling

**DOI:** 10.1101/2020.03.28.013573

**Authors:** Nikhil Maroli

## Abstract

T-2 toxin is one of the potent mycotoxins responsible for several pathological conditions in humans and animals. This nonvolatile low molecular weight toxin compound shows strong resistance to degradation at room temperature as well as at autoclaving temperature. The inhibition of acetylcholinesterase and cytochrome P450 receptor by T-2 toxin leads to many pathological conditions as these two receptors involved in many pathways. Here, we have used melatonin as a protection agent against T-2 toxin in both receptor systems. Using molecular docking, molecular dynamics simulation and free energy calculations we have shown that melatonin can act as a potent inhibitor of T-2 toxin in acetylcholinesterase and cytochrome P450.

## Introduction

T-2 toxin is one of the mycotoxin and bio-warfare agents produced by various species of Fusarium, mainly by *F. sporotrichioides, F. poae, F. acuinatum,* and *F. langsethiae* [1–3]. T-2 toxin commonly presents in cereal grains, including rice, wheat, maize, oats and barley and also presents in cereal-based processed food products. A high concentration of nonvolatile toxins can occur in agriculture products. This low-molecular-weight toxic compound (MW = 466 g/mol) with strong resistance to degradation at room temperature as well as for autoclaving temperatures. The presence of T-2 toxin in agriculture products makes them a potential threat to animals and human. The trichothecenes having a tetracyclic sesquiterpenoid 12,13-polytrichoses-9-one ring and the 12, 13-epoxy ring responsible for the toxicity. The T-2 and HT-2 toxin belongs to type A trichothecenes class, in which carbonyl groups at C8 position are not present. The type B having a carbonyl group at C8 position and in type C, a second epoxy ring between C-7 and C-8 are present. Whereas, type D contains a macrocyclic ring between C-4 and C-15. T-2 toxin is nonvolatile and resistant to degradation in normal environmental conditions such as temperature and light but it is degradable in strong acid or alkaline conditions [4].

The T-2 toxin leads to several pathological conditions such as weight loss, anemia, a decrease in leukocyte count and other related toxicity in the liver and stomach. Recent studies show that the T-2 toxin leads to alimentary toxic aleukia (ATA) and Kashin-Beck disease (KBD)-a chronic endemic osteochondropathy [5–9]. Further, various studies in animals revealed the chronic toxic nature of the T-2 toxin such as cerebral edema, nephrogenic diabetes insipidus. Acute injection of T-2 toxin in rat shows feed refusal and histopathological changes in the liver, spleen, and brain. Systematic injection of T-2 toxin shows the toxic effects such as catarrhal gastritis, necrosis of the lymphoid in rabbit and marginal differences in egg production and impaired egg hatch observed in case of chickens. The long-term application of T-2 toxin in low dosage to mice resulted in the productions of papillomae and carcinomas. The exposure of T-2 toxin results in leukopenia and cell depletion in lymphoid organs and inhibits the erythropoiesis in the bone marrow and spleen. It is reported that the T-2 toxin inhibits IL-2 and IL-5 production by T-cells and it also targets hematopoietic progenitors such as granulocyte, monocyte, and erythrocyte; in results, T-2 toxin attacks the immunity system and which attracts many other pathogens. In addition to that, the T-2 toxin is a well-known protein, DNA and RNA inhibitor which also interacts with lipid bilayers and increases lipid peroxides using the metabolism in the membrane phospholipids. Consequently, the interaction of phospholipids by T-2 toxin generates free radicals, which damage the plasma membrane and lead to cell death. T-2 toxin-induced cell cycle alteration was revealed through the mitogen-activated protein kinase pathway and apoptosis in human neuroblastoma cells [10]. Our studies revealed the aquaporin-4 mediated cerebral edema through T-2 toxin-induced apoptosis and the effect on the renal system through aquaporin-2 [11–12]. T-2 toxin generates pro-apoptotic conditions by initiating the cascade reactions and Fs up-regulation of p53 proteins which in turn increases the Bax/Bcl-2 and Bax/BclxL reactions. Also, JNK/p38 MAP kinase and ERK1/2 pathway activated by stress responses play a crucial role in apoptosis by the T-2 toxin. Studies on rat revealed that the toxicity of the T-2 toxin induces apoptosis in basal keratinocytes which pass through the embryo. The metabolism of T-2 toxin occurs mainly in the liver and the metabolites of T-2 are an HT-2 toxin, T-2 tetraol, T-2 triol, 30-hydroxy HT-2, 30-hydroxy T-2 triol, neosolaniol, 30-hydroxy T-2, and dihydroxy HT-2 and de-epoxy-30-hydroxy T-2 and diepoxy-30-hydroxy HT-2 [2]. The protein and DNA synthesis are inhibited by thiol groups and it also causes oxidative damage to the cells. The inhibition of peptidyl-transferase at the transcription site by T-2 toxin inhibits protein synthesis and it prevents polypeptide chain initiation by inhibiting the 60S ribosomal unit. Furthermore, it inhibits DNA polymerases, monoamine oxidase, terminal deoxynucleotidyl transferase, and many more proteins

Several compounds that show antioxidant properties are widely used to treat against the T-2 toxin such as vitamins, carotenoids, provitamins, chlorophyll, and its derivatives. Recently researchers focus on melatonin against toxin-induced apoptosis and oxidative stress. Melatonin is a hormone secreted by the pineal gland shown to be present in the gut, lymphocytes, bone marrow cells. Furthermore, the highly lipophilic and hydrophilic nature of melatonin made it access the cells and subcellular compartments including the nucleus. Being a highly beneficial molecule, melatonin has significantly been explored in many pathophysiological conditions including inflammation, hepatotoxicity, cancer, cardiovascular disorders, neurological disorders. Earlier reports show that the melatonin act as a potent inhibitor for various mycotoxins such as Ochratoxin A, fumonisin B1 and aflatoxin B1. Here, we first-time study has been designed to elucidate the molecular mechanisms of T-2 toxin inhibition by melatonin through Acetylcholinesterase and Cytochrome P450 pathway. Molecular docking was performed to predict the binding locations of the targeted proteins and to create a receptor-ligand complex for MD simulations. 100 ns Molecular dynamics simulation were performed in water to assess the structural stability and dynamics of the proteins in the presence of T-2 toxin and inhibitor molecule. Umbrella simulations and MMPBSA calculations were performed to study the binding energy and interactions of T-2 toxin and melatonin with Acetylcholinesterase and Cytochrome P450.

## Materials and methods

### Molecular Docking and Molecular Dynamics Simulations

The crystal structures of Human Acetylcholinesterase and Cytochrome P450 were retrieved from the RCSB database (www.rcsb.org) with corresponding entry code 4BDT and 1W0E [13–14], the small molecules melatonin and T-2 toxins were obtained from PubChem compound database. The missing residues were added using the program GalaxyFill [14] after removing the coexisting ligand molecules in the PDB file. In the meantime, melatonin and T-2 mycotoxin structures were geometrically optimized with Gaussian 09 using the B3LYP of density functional theory and 6-31G (d,p) basis set. Docking of ligand molecules with receptors was carried out using Autodock Vina [16]. The docked studies were performed to investigate the interaction of molecules with proteins, further to create receptor-ligands complexes to conduct molecular dynamics simulations. The ligand topology files were prepared using the *cgenff.paramchem.org* server [17–18] and the receptor-ligand model used for molecular dynamics simulation with GROMACS 5.1.2 [19] software package. The LINCS algorithm [20] was used to constrain all covalent bonds between the hydrogen and heavy atoms. The Particle Mesh Ewald method [21] was used for long-range non-bonded interaction with appropriate cutoff length. A leap-frog algorithm was used to integrate the equation of motion of the system, TIP3P water model [22] and all-atom Charmm36 [23] force field was used for all the simulations. Sodium Chloride ions were added in the most favorable position to compensate for the net charge of the receptors. The resulting system comprised of proteins, ligands, water molecules and neutral ions, which is further minimized for 50000 steps and energy convergence attained. The equilibration in NVT and NPT ensemble conducted allowing an integration step of 2 fs. Velocity rescaling methods were used to maintain the constant temperature at 310 K for NVT equilibration and to simulate isothermal-isobaric ensemble (NPT), the temperature (310 K) and the pressure at 1 bar controlled by individual velocity rescaling and Parrinello-Rahman method. The unrestrained simulation length was 100 ns with a time step of 2 fs and trajectories collected in all 10 ps intervals, the same procedure applied to all the simulations, for all the visualization VMD [24], UCSF Chimera [25] and Pymol [26] were used.

### Free energy calculation-Umbrella simulation

The binding free energy of receptor-ligand molecules was calculated using umbrella simulation. To generate equilibrated starting structure for the pulling simulations protein-bound complex placed in a box of a simple point charge (SPC) water [27], and an appropriate number of ions were added to neutralize the system. The small molecules where pulled along the z-direction and box size was maintained to satisfy the minimum image convention. The system was equilibrated at NPT ensemble and temperature was maintained at 310 K using the Berendsen weak coupling method. The protein kept as a reference for pulling simulations and ligands were pulled away from the protein structure along z-axis over 500 ps using a spring constant of 1000 kJmol^-1^nm^-2^ with a pull rate of 0.01 nmps^-1^. Approximately 5 nm was achieved between the center of mass (COM) of protein and ligand. The snapshot was taken to generate starting configurations for umbrella sampling windows. Window spacing (0.2 nm and 0.1 nm) was chosen in a way to ensure sufficient overlapping to obtain a smooth profile and resulted in 46 and 54 windows. In each window, a 10 ns simulation was performed for umbrella sampling. The analysis was carried out with the weighted histogram analysis method (WHAM) [28].

### Free energy calculation-MMPBSA

The umbrella simulation method provides precise binding energy. However, the individual residue contribution towards the binding energy is still unknown. In this regard, we used the molecular mechanics Poisson-Boltzmann surface area (MM/PBSA) method to calculate residue wise contribution and binding energy implemented in g_mmpbsa.

Binding free energy of Protein-Ligand complex in solvent is expressed as

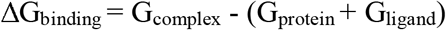

Where G_complex_ is the total free energy of protein-ligand complex, G_pro_t_ei_n and G_ligand_ are total energy of separated protein and ligand in a solvent, respectively.

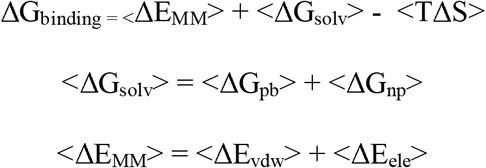

<Δ*E_MM_*> is the total molecular mechanics energy in the gas phase, TΔS is entropy and <ΔG_solv_> is salvation free energy. <ΔE_MM_> is the sum of electrostatic and van der Waals interaction energy.

## Results and Discussion

The present study revealed that melatonin could find potential inhibitory action against T-2 toxin in a mammalian system. We used molecular dynamics simulations to elucidate the role of melatonin as a potent inhibitor of T-2 toxin through AChE and CP450 receptor-mediated pathways. Further, the inhibition mechanism of T-2 toxin was revealed by studying the structural and dynamical changes of the proteins with toxin and inhibitor.

### Molecular docking

The binding modes of AChE and CP450 are different from each other, though there was a significant contribution of residues in both receptors such as Trp, Tyr, Ser in the binding sites [4–8]. Recent studies on the structural aspects of CP450 revealed the importance of hydrogen bonds in the binding site, which also revealed the conformational changes induced by the ligand molecules at the binding site. Figure 1 shows the three-dimensional conformations of docked complexes, AChE with melatonin shows one hydrogen bond of length 2.80 Å with 337Tyr and hydrophobic interaction with eleven amino acids while toxin shows interactions with nine amino acid residues (Figure S1). The LigPlot diagrams of receptor-ligand interactions at the binding sites are provided in the supplementary information (Figure S2). The toxin with CP450 shows two hydrogen bonds with residue 212Arg spanning a length of 2.99 Å and 3.18 Å, whereas melatonin shows only one hydrogen bond of length 3.09 Å with residue 224Thr. The details of docked results are summarised in Table 1.

**Figure 1.**
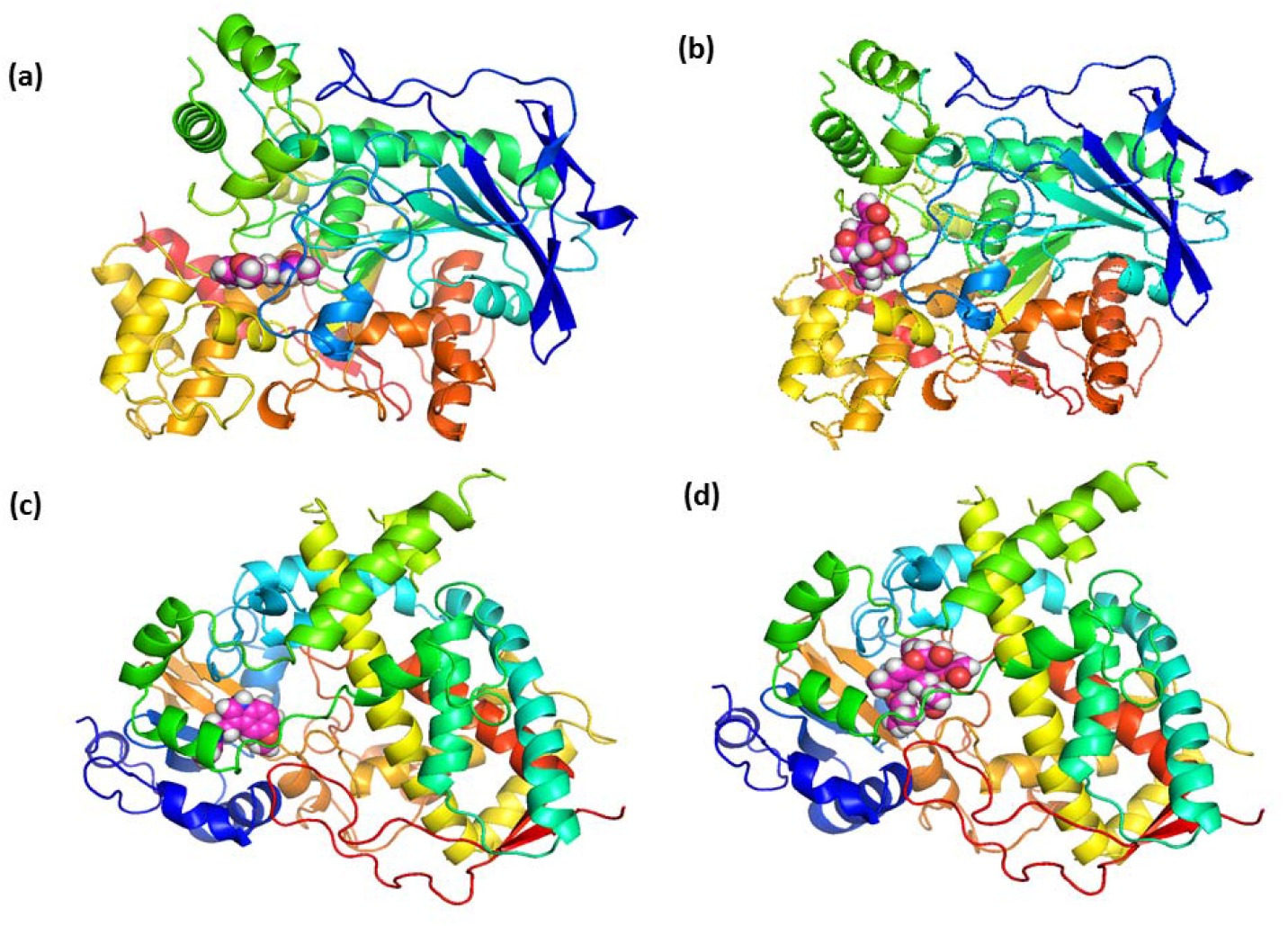
3-D conformations of ligands bound to ACHE and CP450. (a) AChE-melatonin, (b) AChE-T-2 toxin (c) CP450-melatonin (d) CP450-T-2 toxin.

**Table 1.** The affinity value of ligands with proteins and interacting residue for Hydrogen bond and Hydrophobic interactions.(interaction energies are given in kcal/mol for conveniance)

| Protein<br>s | Ligand's | Affinity<br>(kcal/mol<br>) | No. of H-Bonds | H-Bonding Interacting<br>Residue | Hydrophobic Interacting<br>residue |
| --- | --- | --- | --- | --- | --- |
| AChE | Melatonin | -7.6 | 1 | Tyr 337 | Val294,Tyr341,Phe295,Phe338,Phe297,Gly122,Tyr124,Gly121,Tyr72,Tyr337,Trp286 |
|  | T2-toxin | -6.4 | 0 |  | Tyr341, Val294, Ser293, Tyr72, Asp74, His287, Asn283, Trp 286,Leu76 |
| CP450 | Melatonin | -7.9 | 1 | Thr224 | Met371,Arg106,Phe215,Arg372,Phe57,Asp76,Tyr53,Leu216,Leu221,Thr224 |
|  | t2-toxin | -7.7 | 1 | Arg212 | Phe108,Ala370,Arg106,Phe57,Gly481,Arg372,Glu374,Ser119,Arg105,Phe215,Met371,Arg212 |

### Molecular Dynamics (MD) Studies

To assess the convergence and stability of the proteins complex the root mean square deviations (RMSDs) and root mean square fluctuations (RMSFs) of backbone atoms of the proteins were obtained (Figure S3). The RMSD and RMSF indicate that the melatonin system reAChEs equilibration earlier and the average backbone RMSD with melatonin was 0.15 nm and 0.16 nm with AChE and CP450, respectively. Whereas the toxin bound systems show slightly higher fluctuations of 0.16 nm and 0.17 nm, respectively. The non-bonded interaction energies including hydrogen bonds were calculated to understand the various interaction and their contribution to the stability of the inhibitors. The hydrogen bonds between the amino acid residues of the receptors and ligand molecules were calculated by applying the cutoff criteria; distance ≤ 3.0 Å; angle ≥ 120 degrees[29–30]. The electrostatic and van der Waals interaction energy has a dominant role in protein-ligand interactions. The average non-bonded interaction energy and the nature of the interactions are in favor of the melatonin bound receptors, whereas toxin bound receptors show unstable and higher energy. It can be noted that the interaction energies are given on a negative scale (Figure 2). The binding cavity of AChE is mainly hydrophobic because of the existence of several non-polar amino acids (Trp286, Val294, Phe338, Phe295, Phe297, Gly121, and Gly122). It is also noted that the presence of pi-pi stacking between the melatonin and amino acid residues such as Trp286, Tyr124 and Trp439 and pi-alkyl interaction between Val294 in the binding cavity. Recent studies also showed that the pi-pi stacking interaction with the aromatic and indole ring of Trp residues at the binding site of the AChE and CP450 receptor. We also observed strong non-bonded interactions in AChE bound systems. The AChE toxin bound system shows pi-pi stacking between the toxin and residues Trp286, Tyr72 and pi-alkyl interaction with Tyr341 residue. The average van der Waals interaction energy between the melatonin is higher than the toxin bound system. The presence of non-polar amino acids in the binding cavity of AChE stabilizes the inhibitors via van der Waals interaction energy; in effect, both electrostatic and van der Waals energies are favors towards the stability of the inhibitory molecule. The binding site of CP450 contains both non-polar and polar amino acids (Met371, Arg106, Phe215, Thr224, Leu216, Leu221, Tyr53, Asp76, Phe57, and Arg372). Pi-Pi stacked interaction observed between melatonin and residues such as Phe220 and Phe215 and the presence of pi-alkyl interactions was observed with Tyr341, Leu76, His287, and melatonin. In toxin bound system pi-alkyl interactions observed with residues Phe108, Phe215, Arg372, Met371, and Phe57 (Figure S5). As a result, the presence of both polar and non-polar amino acids in the binding site contributes to higher non-bonded interaction energy in CP450 protein systems. Further, the secondary structural changes and the radius of gyration of the proteins were calculated from the MD trajectories to investigate the effect of the molecules on receptors (Figure 3). AChE bound to melatonin shows a few sequences (60-100) of residues changed to turn and bend from Alpha helix and there are no noticeable structural changes from sheet to the helix or vice versa. The AChE possesses 32.6% helix, 16.7% sheet, 14.4% turn, 33.9% coil and 2.5% 3-10 helix in their secondary structure. The melatonin bound AChE shows 32.3% helix, 16.3% sheet, 15.4% turn, and 35.9% coil at the end of the simulations. Whereas toxin bound, AChE shows more transitions in secondary structure, which consists of 32.3% helix, 15.5% sheets, 11.0%, turn, 40.5% coil and 0.8% 3-10 helix at the end of the simulation. This depicts the conformational changes occurs in T-2 toxin bound proteins. The post docked CP450 shows 46.1% helix, 8.7% sheet, 10.4% turn and 34.8% coil. At the end of the 100 ns simulation CP450 bound with melatonin shows 46.1% helix, 8.9% sheet, 12.2% turn, 32.0% coil and 0.9% 3-10 helix, whereas toxin bound system shows 49.3% helix, 8.9% sheet, 10.4% turn,30.4% coil and 0.9% 3-10 helix.

**Figure 2.**
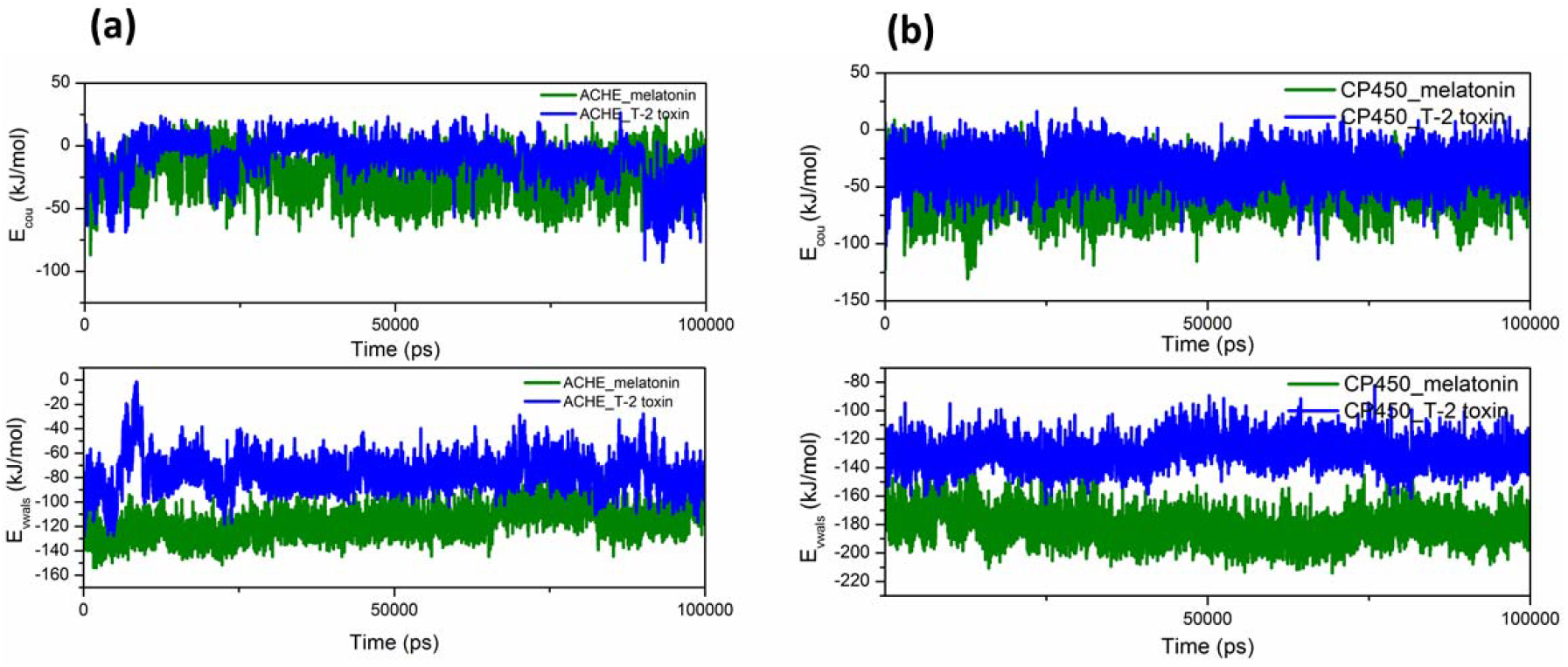
The electrostatic and van der Waals interaction energy between the receptors and ligands. (a) AChE bound to melatonin and T-2 toxin (b) CP450 bound to melatonin and T-2 toxin.

### Principal Component Analysis (PCA)

To understand the collective motion of the receptor, we performed PCA based on Cα atoms in the protein denoted by the eigenvectors of the covariance matrix [31]. The Root means square fluctuations of Cα atoms along the first two eigenvectors are shown in Figure 3. The deviation of RMSF along the two eigenvectors corresponding to the structural changes in the protein. The root means square fluctuations of AChE-melatonin bound systems show collective fluctuations along both vectors corresponding to the residues 68-78, which shows minor changes in the secondary structural analysis. AChE-toxin bound system shows higher fluctuations of RMSF corresponding to the residues 53-68, 19-21, and 76-98. The higher deviations in the RMSF might be due to the occasional deviations from helix to random coil or bends, which is again consistent with secondary structural analysis. The bound melatonin system with CP450 shows fluctuations corresponding to the residues 60-70 and 110-120. The toxin bound CP450 shows higher fluctuations for the residues 60-70 than melatonin bound systems. Melatonin bound proteins show fewer deviations in RMS values depicts enhanced structural stability.

**Figure 3.**
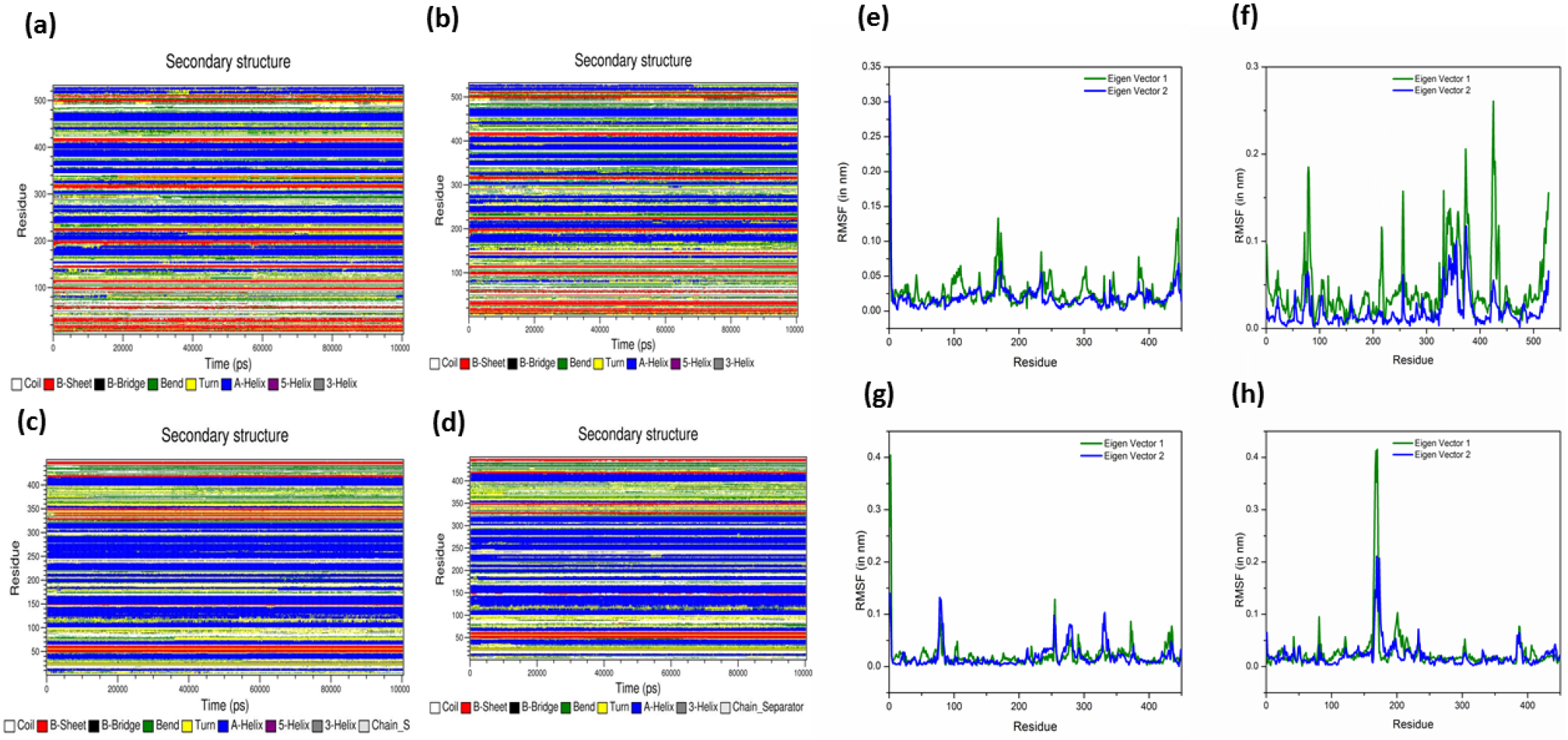
Time evolution of secondary structure and RMSF along First two eigen vectors. (a) & AChE with melatonin (b) AChE with T-2 toxin (c) CP450 with melatonin (d) CP450 with T-2 toxin

To understand the possible structural conformations and stability the free energy landscape of protein-bound complex obtained based on the radius of gyration and root means square deviation. The Free energy landscape of all the systems is provided in Figure S7 of supplementary information. The energy landscape analysis shows that the bound melatonin system achieved faster and stable conformation than the toxin bound systems. The local minima or global minima is connected with the energy barrier, and the AChE-toxin bound system shows multiple local minima whereas melatonin shows two lower energy basins. CP450 bound with toxin shows multiple minima corresponding to meta-states whereas melatonin shows two minima. In both systems, toxin possesses multiple minima connected with energy barriers whereas melatonin shows comparatively stable states of the system.

### Free energy calculations

#### Umbrella Simulation

The binding energy ΔG, calculated using the umbrella simulation method along the reaction coordinate. Potential of mean force (PMF) curves were obtained for each molecule (Figure 4), leading to ΔG of binding for melatonin and T-2 toxin. A better PMF can be achieved by extending the simulation time and adding more umbrella windows, but the present calculation is sufficient to demonstrate our aim. The PMF in both proteins dramatically increased in a range of 0 nm to 2.2 and 3. 5 nm. Afterward, the slope of the PMF became mild. The ΔG is calculated by the difference between the plateau region of the PMF curve and the energy minimum of the curve. In both receptors systems, melatonin shows higher PMF values of 8.74 kJ/mol and 17.42 kJ/mol respectively, while the value of T-2 toxin is 5.09 kJ/mol and 10.67 kJ/mol respectively. These results indicate the interactions between the receptors and the T-2 toxin is weaker than the melatonin, which is consistent with previous analysis. The umbrella sampling results confirmed that the melatonin possesses higher binding energy than T-2 toxin in both the protein system.

**Figure 4.**
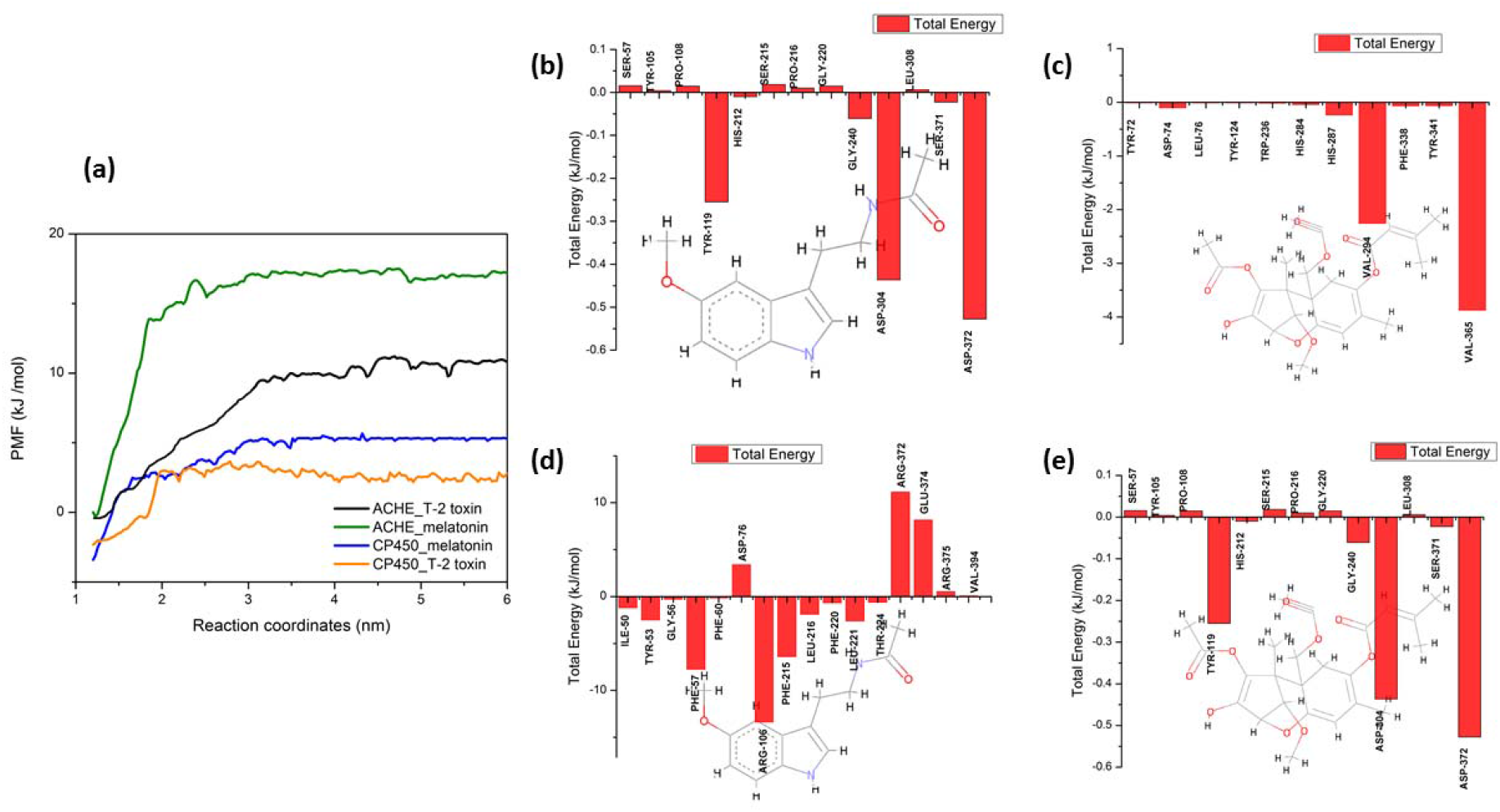
(a) Potential of mean force (PMF) of melatonin and T-2 toxin with AChE and CP450 receptors. (b) and (c) represents the residue wise contribution at the binding site for ACHE-melatonin and AChE -T-2 toxin. (d) and (e) represents the residue wise contribution at the binding site for CP450-melatonin and CP450-T-2 toxin

### MM-PBSA

MM/PBSA method was used to understand the individual contribution of residues towards the binding energy and contributions from different energy components. The last 10 ns frames were extracted from the 100 ns simulation and calculations are performed using g_mmpbsa. The binding energy and the contribution from electrostatic, van der Waals and polar solvation energies are summarised in Table 2. In both systems, the melatonin bound complex shows higher binding energy. The per-residue contribution of binding energy in the active sites of the proteins is given in Figure 4. In all model systems, van der Waals interaction energy contributes negatively to the binding energy whereas polar solvation energy contributes positively towards the binding energy.

**Table 2.**
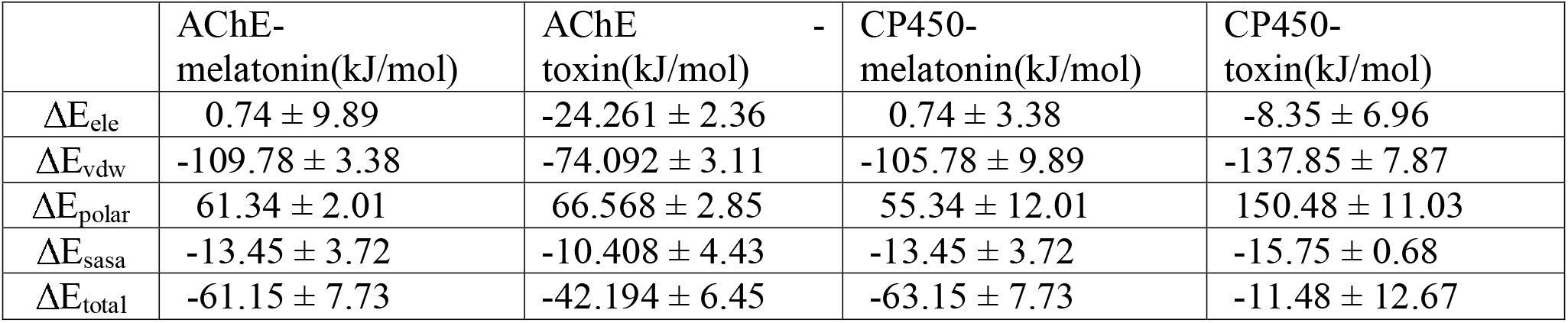
The binding free energy and different contribution towards the binding energy in AChE and CP450 protein system.

## Conclusion

We have shown the binding mechanism of melatonin and T-2 toxin and how melatonin can be used to protect the T-2 toxin led toxicity via AChE and CP450 mediated pathways. The post-processing MD analysis including PCA and secondary structural changes of receptors with melatonin and T-2 toxin shows that the presence of T-2 toxin leading to conformational transition. The electrostatic and van der Waals interaction energies indicate that the presence of non-polar amino acids in the AChE binding site plays a major role in the stability of melatonin, whereas in CP450 presence of both polar and non-polar amino acids stabilize the melatonin. The PCA analysis shows the higher fluctuations in the presence of T-2 toxin over melatonin indicates the stability of proteins in the presence of melatonin. The results obtained from PCA and MD trajectory analyses highlighted the structural and dynamical changes occur in the presence of t2-toxin. Umbrella sampling simulation and MM-PBSA calculations also depict higher binding energy of melatonin over T-2 toxin. The interaction of T-2 toxin in both receptors is weaker than the melatonin. However, melatonin does not lead to any conformational changes to the receptors which indicated that the melatonin can be used as a potential inhibitor for the T-2 toxin.

## References

[1] Sudakin, D. L. (2003). Trichothecenes in the environment: relevance to human health. Toxicology letters, 143(2), 97–107.

[2] Mateo, J. J., Mateo, R., & Jimenez, M. (2002). Accumulation of type A trichothecenes in maize, wheat and rice by Fusarium sporotrichioides isolates under diverse culture conditions. International Journal of Food Microbiology, 72(1-2), 115–123.

[3] D’mello, J. P. F., Placinta, C. M., & Macdonald, A. M. C. (1999). Fusarium mycotoxins: a review of global implications for animal health, welfare and productivity. Animal feed science and technology, 80(3-4), 183–205.

[4] Li, Y., Wang, Z., Beier, R. C., Shen, J., Smet, D. D., De Saeger, S., & Zhang, S. (2011). T-2 toxin, a trichothecene mycotoxin: review of toxicity, metabolism, and analytical methods. Journal of agricultural and food chemistry, 59(8), 3441–3453.

[5] Ishigami, Noriaki, and Shinya Sehata. “T-2 toxin-induced toxicity in pregnant mice and rats.” International journal of molecular sciences 9.11 (2008): 2146–2158.

[6] Čonková, E., Laciakova, A., Kováč, G., & Seidel, H. (2003). Fusarial toxins and their role in animal diseases. The veterinary journal, 165(3), 214–220.

[7] Rafai, P., Bata, A., Vanyi, A., Papp, Z., Brydl, E., Jakab, L., … & Tury, E. (1995). Effect of various levels of T-2 toxin on the clinical status, performance and metabolism of growing pigs. The Veterinary Record, 136(19), 485–489.

[8] MacDonald, E. J., Cavan, K. R., & Smith, T. K. (1988). Effect of acute oral doses of T-2 toxin on tissue concentrations of biogenic amines in the rat. Journal of animal science, 66(2), 434–441.

[9] Shinozuka, J., Suzuki, M., Noguchi, N., Sugimoto, T., Uetsuka, K., Nakayama, H., & Doi, K. (1998). T-2 toxin-induced apoptosis in hematopoietic tissues of mice. Toxicologic pathology, 26(5), 674–681.

[10] Agrawal, M., Bhaskar, A. S. B., & Rao, P. L. (2015). Involvement of mitogen-activated protein kinase pathway in T-2 toxin-induced cell cycle alteration and apoptosis in human neuroblastoma cells. Molecular neurobiology, 51(3), 1379–1394.

[11] Maroli, N., Jayakrishnan, A., Ramalingam Manoharan, R., Kolandaivel, P., & Krishna, K. (2019). Combined Inhibitory Effects of Citrinin, Ochratoxin-A, and T-2 Toxin on Aquaporin-2. The Journal of Physical Chemistry B, 123(27), 5755–5768.

[12] Maroli, N., Kalagatur, N. K., Bhasuran, B., Jayakrishnan, A., Manoharan, R. R., Kolandaivel, P., … & Kadirvelu, K. (2019). Molecular Mechanism of T-2 Toxin-Induced Cerebral Edema by Aquaporin-4 Blocking and Permeation. Journal of chemical information and modeling, 59(11), 4942–4958.

[13] Nachon, F., Carletti, E., Ronco, C., Trovaslet, M., Nicolet, Y., Jean, L., & Renard, P. Y. (2013). Crystal structures of human cholinesterases in complex with huprine W and tacrine: elements of specificity for anti-Alzheimer’s drugs targeting acetyl-and butyryl-cholinesterase. Biochemical Journal, 453(3), 393–399.

[14] Williams, P. A., Cosme, J., Vinković, D. M., Ward, A., Angove, H. C., Day, P. J., … & Jhoti, H. (2004). Crystal structures of human cytochrome P450 3A4 bound to metyrapone and progesterone. Science, 305(5684), 683–686.

[15] Coutsias, E. A., Seok, C., Jacobson, M. P., & Dill, K. A. (2004). A kinematic view of loop closure. Journal of computational chemistry, 25(4), 510–528.

[16] Trott, O., & Olson, A. J. (2010). AutoDock Vina: improving the speed and accuracy of docking with a new scoring function, efficient optimization, and multithreading. Journal of computational chemistry, 31(2), 455–461.

[17] Vanommeslaeghe, K., Hatcher, E., Acharya, C., Kundu, S., Zhong, S., Shim, J., … & MacKerell Jr, A. D. (2010). CHARMM general force field: A force field for druglike molecules compatible with the CHARMM all-atom additive biological force fields. Journal of computational chemistry, 31(4), 671–690.

[18] Yu, W., He, X., Vanommeslaeghe, K., & MacKerell Jr, A. D. (2012). Extension of the CHARMM general force field to sulfonyl containing compounds and its utility in biomolecular simulations. Journal of computational chemistry, 33(31), 2451–2468.

[19] Hess, B., Kutzner, C., Van Der Spoel, D., & Lindahl, E. (2008). GROMACS 4: algorithms for highly efficient, load-balanced, and scalable molecular simulation. Journal of chemical theory and computation, 4(3), 435–447.

[20] Hess B, Bekker H, Berendsen HJC, Fraaije JGEM. LINCS: A linear constraint solver for molecular simulations. Journal of Computational Chemistry. 1997; 18:1463–1472.

[21] Darden T, York D, Pedersen L. Particle mesh Ewald: An N·log(N) method for Ewald sums in large systems. The Journal of Chemical Physics. 1993; 98:10089.

[22] Jorgensen WL, Chandrasekhar J, Madura JD, Impey RW, Klein ML. Comparison of simple potential functions for simulating liquid water. The Journal of Chemical Physics. 1983; 79:926.

[23] Mackerel AD, Bashford D, Bellott M, et al. All-atom empirical potential for molecular modeling and dynamics studies of proteins †. The Journal of Physical Chemistry B. 1998; 102:3586–3616.

[24] Humphrey W, Dalke A, Schulten K. VMD: Visual molecular dynamics. Journal of Molecular Graphics. 1996; 14:33–38.

[25] Pettersen EF, Goddard TD, Huang CC, et al. UCSF Chimera?A visualization system for exploratory research and analysis. Journal of Computational Chemistry. 2004; 25:1605–1612.

[26] Schrödinger, L. “The PyMOL Molecular Graphics System. 1.4.: Schrödinger.”

[27] Berendsen HJC, Postma JPM, van Gunsteren WF, Hermans J. Interaction models for water in relation to protein hydration. In: The Jerusalem Symposia on Quantum Chemistry and Biochemistry. Springer Nature; 1981:331–342.

[28] Kumar S, Rosenberg JM, Bouzida D, Swendsen RH, Kollman PA. THE weighted histogram analysis method for free-energy calculations on biomolecules. I. The method. Journal of Computational Chemistry. 1992; 13:1011–1021.

[29] Maroli, N., & Kolandaivel, P. (2018). Structure, stability and water permeation of ([D-Leu-L-Lys-(D-Gln-L-Ala) 3]) cyclic peptide nanotube: A molecular dynamics study. Molecular Simulation, 44(3), 225–235.

[30] Maroli, N., & Kolandaivel, P. (2020). Comparative study of stability and transport of molecules through cyclic peptide nanotube and aquaporin: A molecular dynamics simulation approach. Journal of Biomolecular Structure and Dynamics, 38(1), 186–199.

[31] Amadei A, Linssen ABM, Berendsen HJC. Essential dynamics of proteins. Proteins: Structure, Function, and Genetics. 1993; 17:412–425.

