## supplementary information for "Melatonin Protects T-2 toxin-induced neuronal stress through Acetylcholinesterase & Cytochrome P450 receptor-mediated signaling"

Nikhil Maroli

Computational Biology Division, DRDO BU CLS, Coimbatore-641046, Tamil Nadu, INDIA.

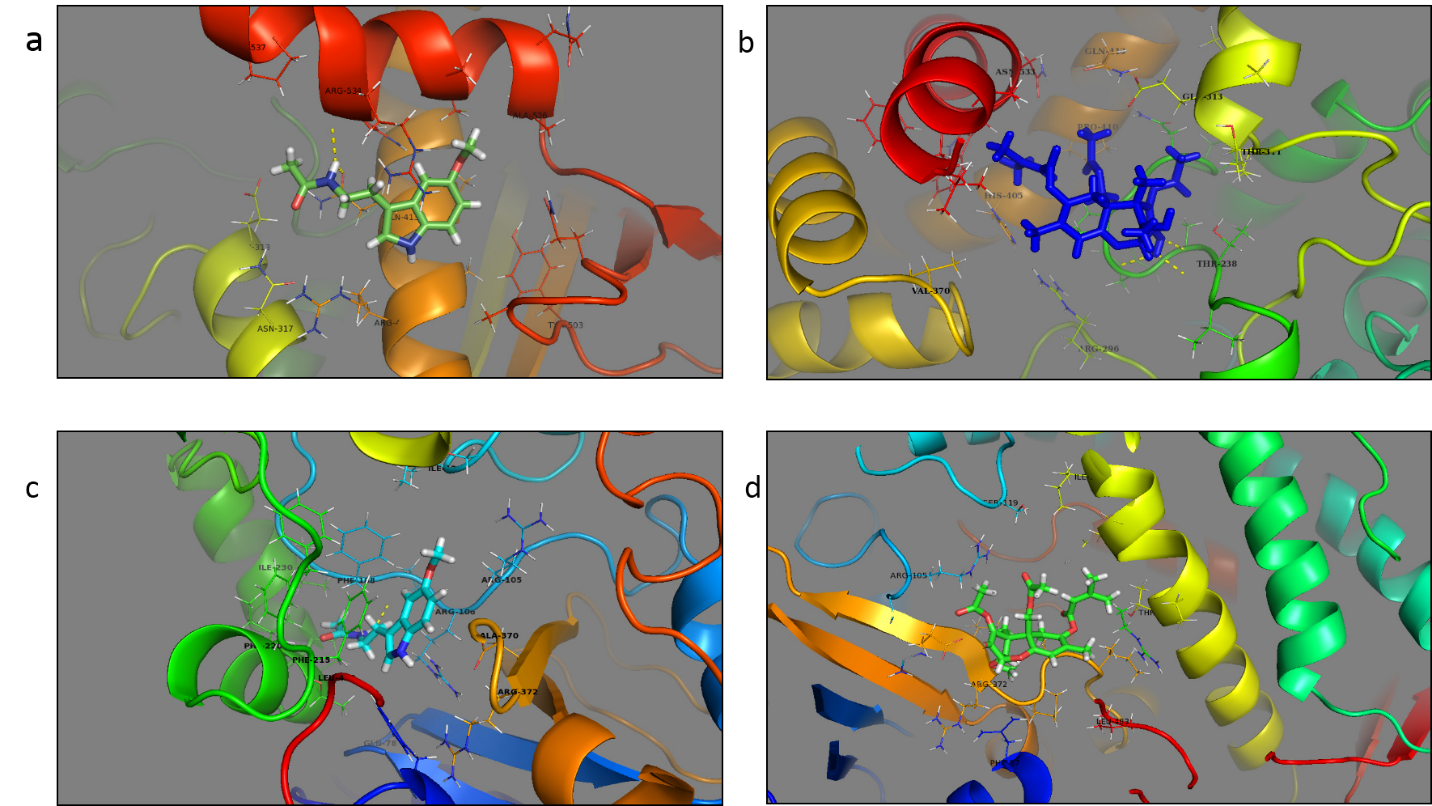

**Figure S1.** The binding site of AChE and CP450, Proteins were represented by New Cartoon models and ligands as stick. (a) AChE-melatonin (b) AChE- T-2 toxin (c) CP450-melatonin (d) CP450- T-2 toxin.

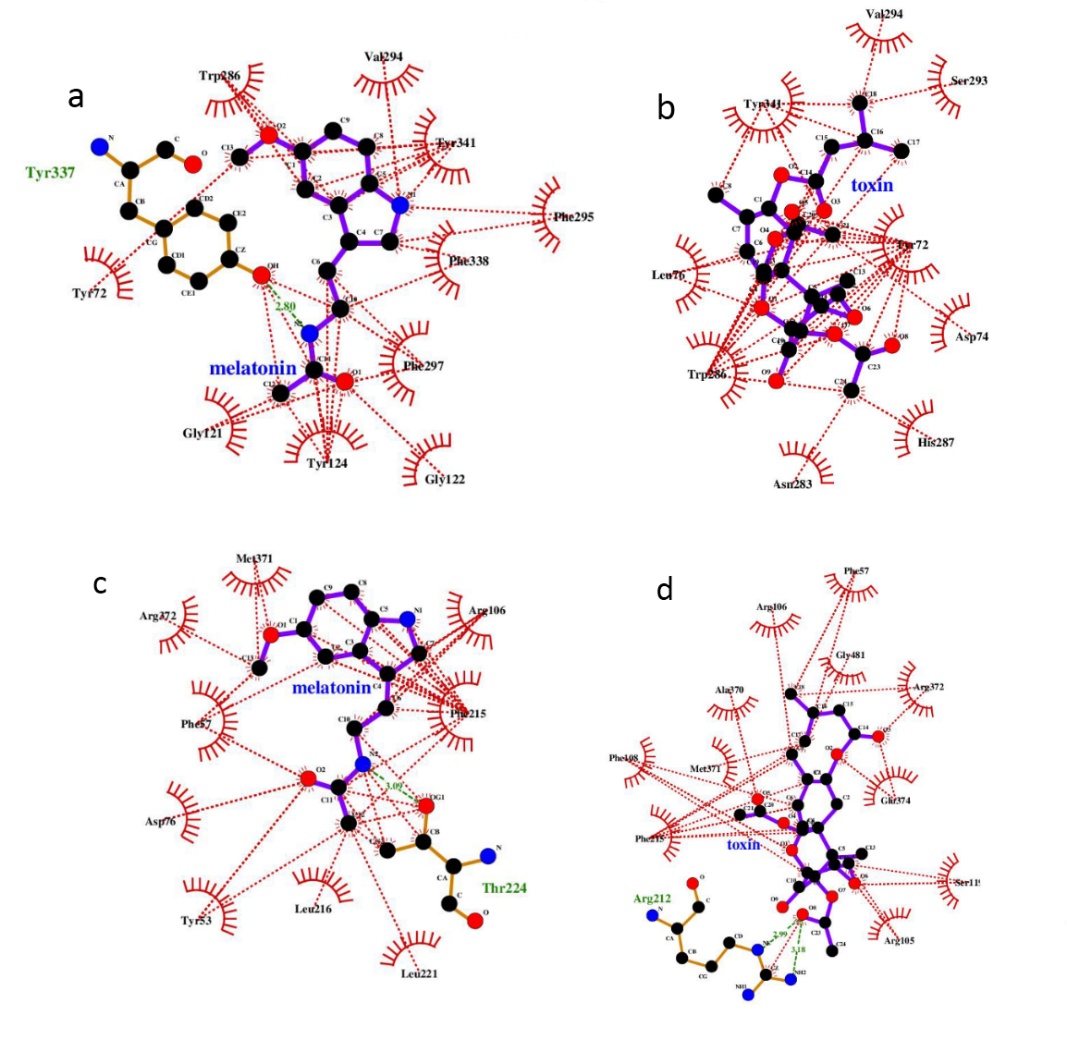

**Figure S2**. Ligplot view of ligands in the binding site, Hydrophobic and hydrogen bonds interactions are represented in red and green dot line respectively. (a) AChE-melatonin (b) AChE-T-2 toxin (c) CP450-melatonin (d) CP450-T-2 toxin.

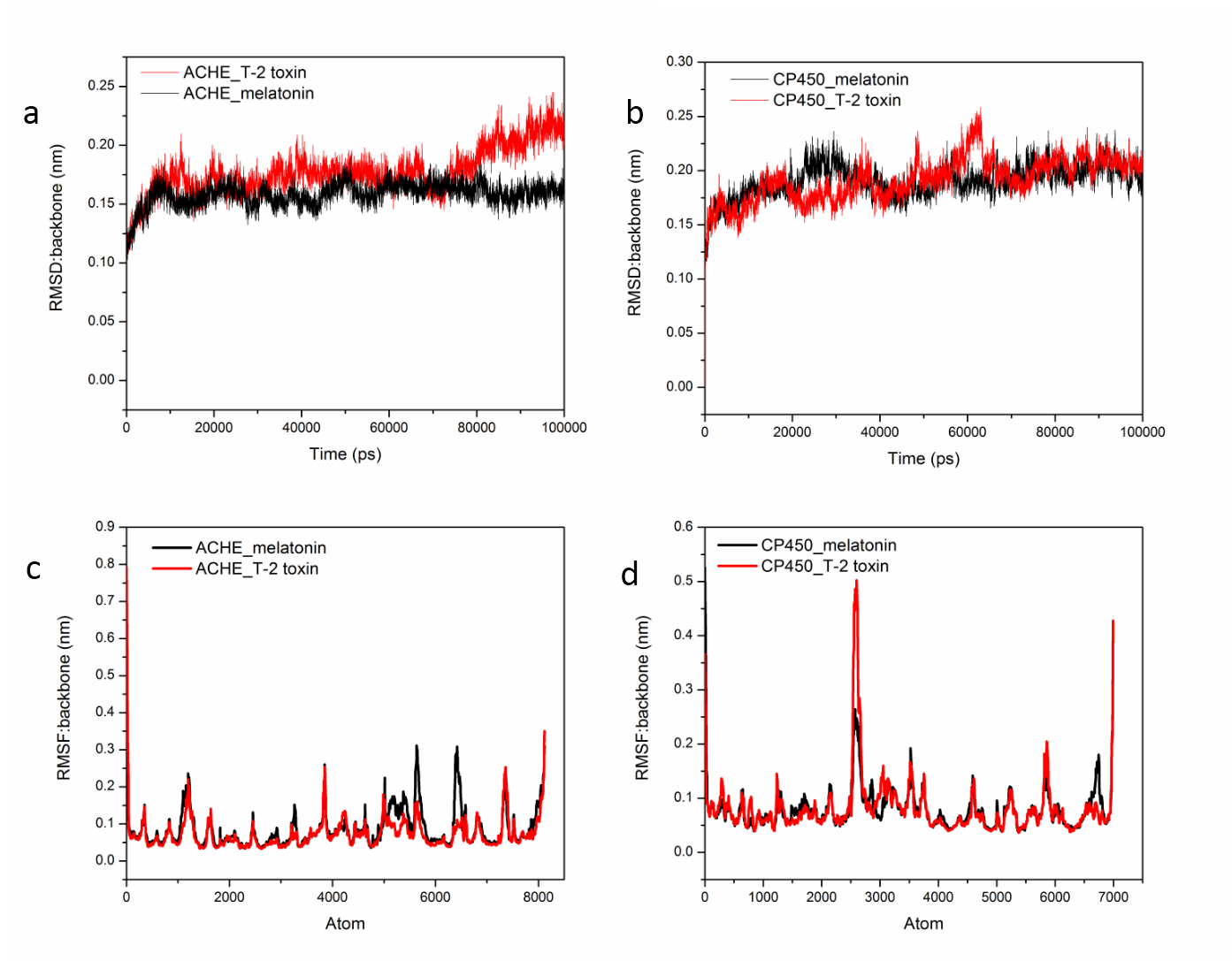

**Figure S3**. Root mean square deviation of backbone atoms and root mean square fluctuations of all atoms of AChE and CP450. (a) RMSD of AChE bound to melatonin and T-2 toxin (b) RMSD of CP450 bound to melatonin and T-2 toxin (c) RMSF of AChE bound to melatonin and T-2 toxin (d) RMSF of CP450 bound to melatonin and T-2 toxin.

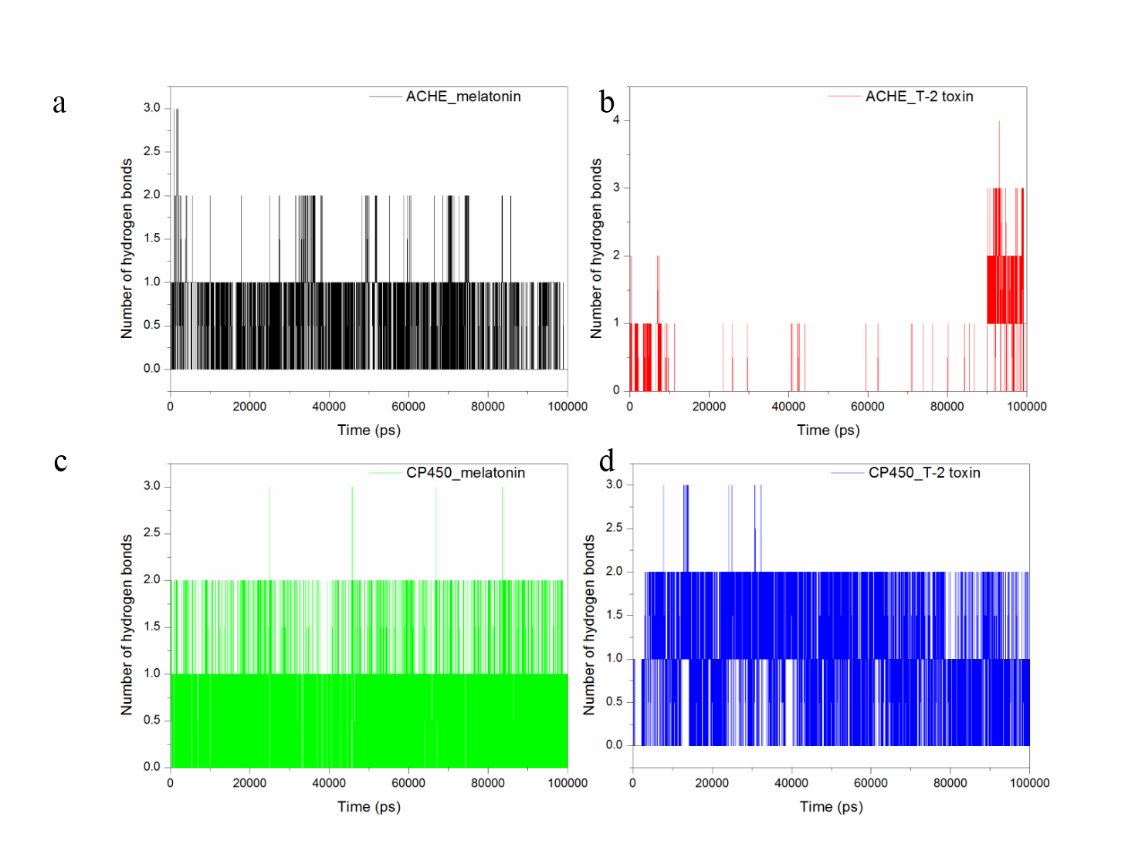

**Figure S4**. The number of hydrogen bonds during the simulation time. (a) AChE bound to melatonin (b) AChE bound to T-2 toxin (c) CP450 bound to melatonin (d) CP450 bound to T-2 toxin.

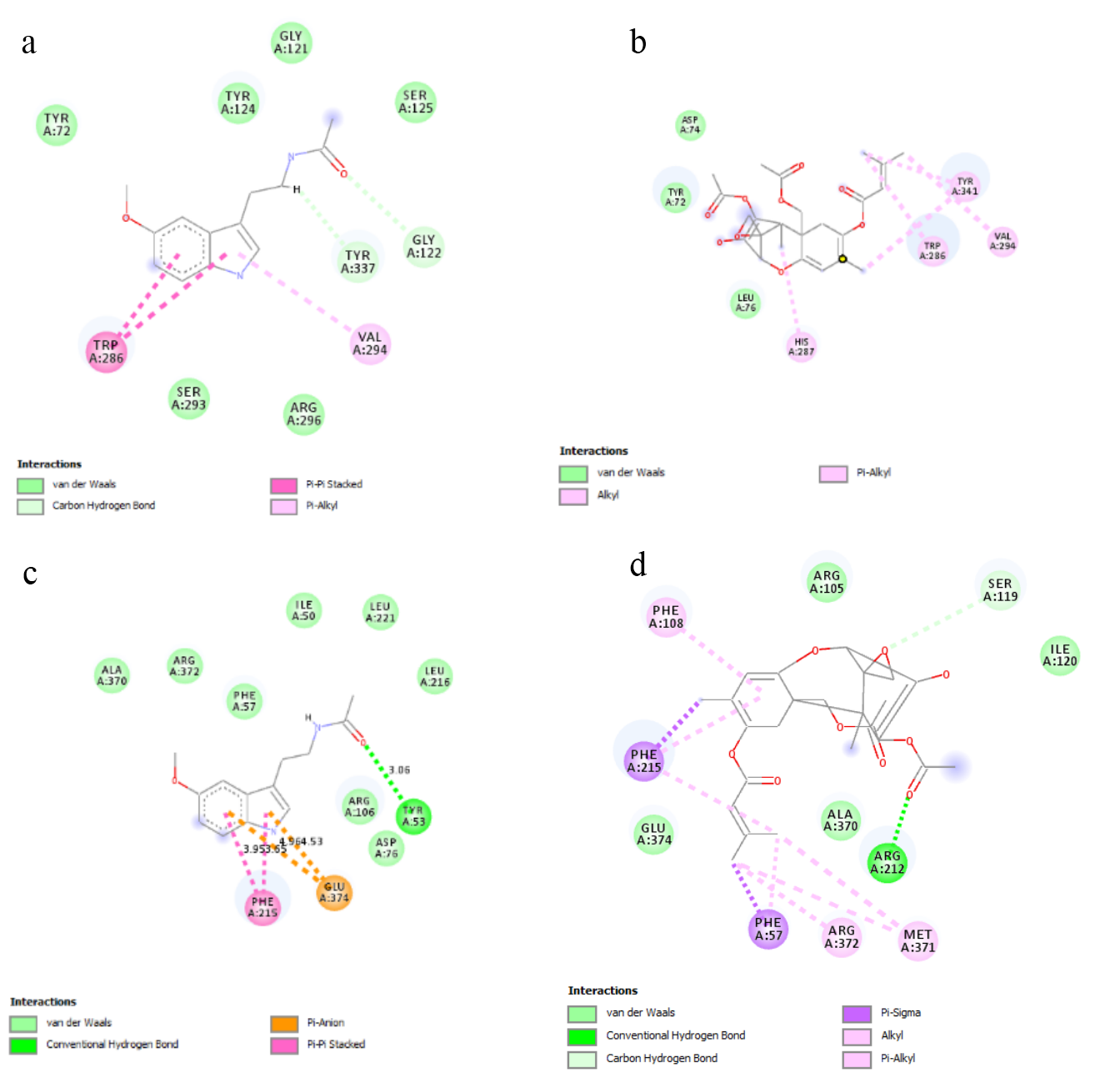

**Figure S5**. 2D diagram of various interactions at the binding site of AChE and CP450. (a) AChE-melatonin (b) AChE- T-2 toxin (c) CP450-melatonin (d) CP450- T-2 toxin.

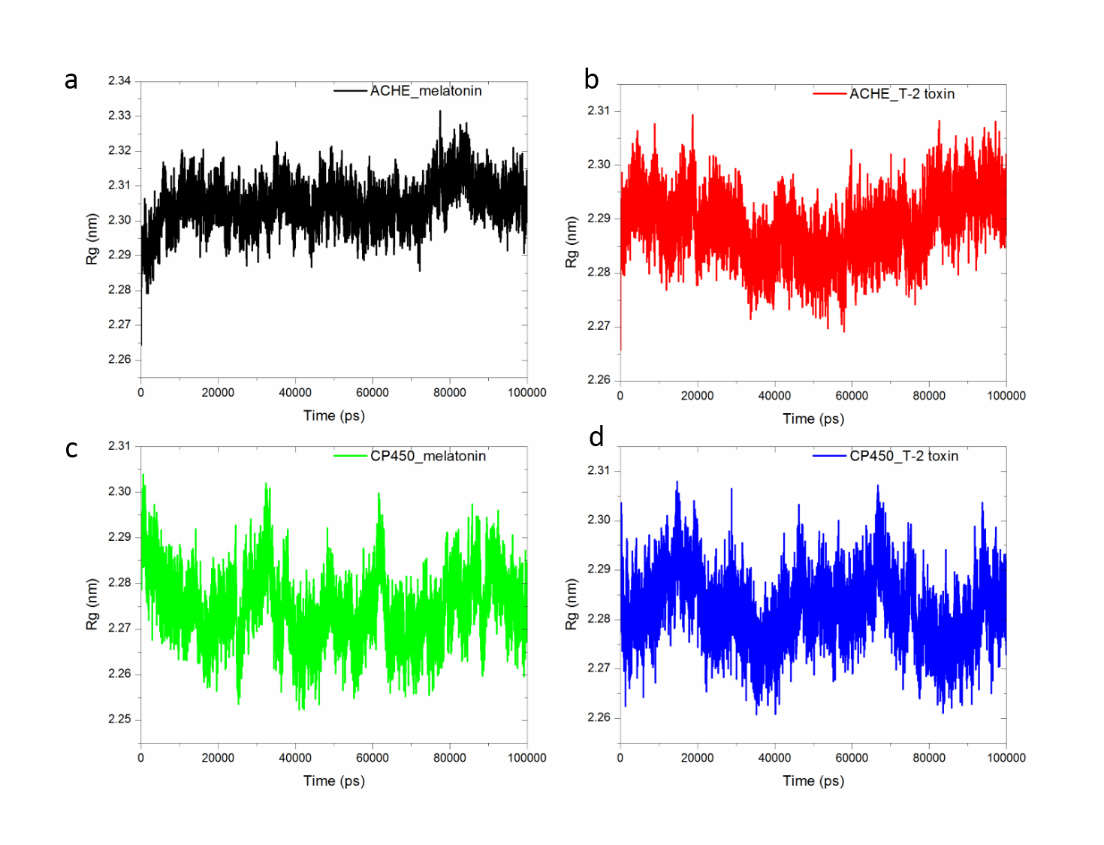

**Figure S6**. The radius of gyration of AChE and CP450 receptors with of melatonin and T-2 toxin. (a) AChE-melatonin (b) AChE-T-2 toxin (c) CP450-melatonin (d) CP450-T-2 toxin.

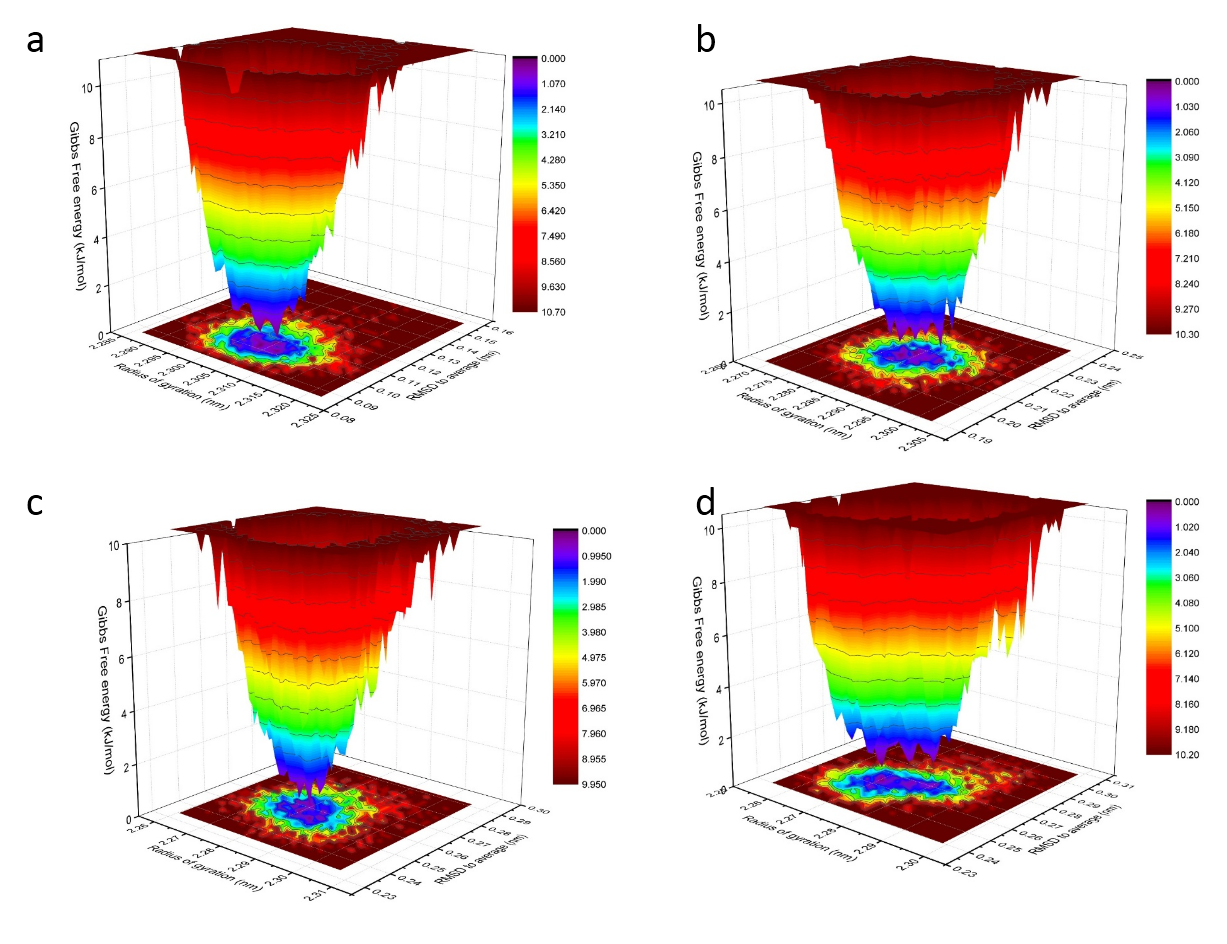

**Figure S7**. Free energy landscape of receptor-ligand system in terms of radius of gyration and root mean square deviations. (a) AChE-melatonin(b) AChE-T-2 toxin (c) CP450-melatonin (d) CP450-T-2 toxin.

| AChE-melatonin | | | |
| --- | --- | --- | --- |
|  | Average | Err.Est | Tot-Drift |
| E_coul_ (kJ/mol) | -23.761 | 2.2 | -4.81 |
| E_vwl_ (kJ/mol) | -117.857 | 3.2 | 20.86 |
| AChE-T-2 toxin | | | |
|  | Average | Err.Est | Tot-Drift |
| E_coul_ (kJ/mol) | -9.83 | 3.7 | -17.69 |
| E_vwl_ (kJ/mol) | -76.74 | 1.2 | -1.36 |

Table 2 Electrostatic and van der Waals interaction energy between AChE-melatonin and AChE-T-2 toxin

| CP450-melatonin | | | |
| --- | --- | --- | --- |
|  | Average | Err.Est | Tot-Drift |
| E_coul_ (kJ/mol) | -50.58 | 1.4 | 3.67 |
| E_vwl_ (kJ/mol) | -182.21 | 1.9 | -8.72 |
| CP450- T-2 toxin | | | |
|  | Average | Err.Est | Tot-Drift |
| E_coul_ (kJ/mol) | -33.8 | 0.86 | 0.129 |
| E_vwl_ (kJ/mol) | -130.08 | 1.8 | 3.01 |

Table 3 Electrostatic and van der Waals interaction energy between CP450-melatonin and AChE-T-2 toxin.
